# Evolutionarily conserved genetic interactions between *nphp-4* and *bbs-5* mutations exacerbate ciliopathy phenotypes

**DOI:** 10.1101/2021.08.25.457729

**Authors:** Melissa R. Bentley-Ford, Melissa LaBonty, Holly R. Thomas, Courtney J. Haycraft, Mikyla Scott, Cameron LaFayette, Mandy J. Croyle, John M. Parant, Bradley K. Yoder

## Abstract

Primary cilia are sensory and signaling hubs with a protein composition that is distinct from the rest of the cell due to the barrier function of the transition zone (TZ) at the base of the cilium. Protein transport across the TZ is mediated in part by the BBSome, and mutations disrupting TZ and BBSome proteins cause human ciliopathy syndromes. Ciliopathies have phenotypic variability even among patients with identical genetic variants, suggesting a role for modifier loci. To identify potential ciliopathy modifiers, we performed a mutagenesis screen on *nphp-4* mutant *C. elegans* and uncovered a novel allele of *bbs-5*. *Nphp-4;bbs-5* double mutant worms have phenotypes not observed in either individual mutant strain. To test whether this genetic interaction is conserved, we also analyzed zebrafish and mice mutants. While *Nphp4* mutant zebrafish appeared overtly normal, *Bbs5* mutants exhibited scoliosis. When combined, *Nphp4*;*Bbs5* double mutant zebrafish did not exhibit synergistic effects, but the lack of a phenotype in *Nphp4* mutants makes interpreting these data difficult. In contrast, viable *Nphp4;Bbs5* double mutant mice were not obtained and there were fewer mice than expected carrying three mutant alleles. Additionally, postnatal loss of *Bbs5* in mice using a conditional allele compromised survival when combined with a *Nphp4* allele. As cilia are formed in the double mutant mice, the exacerbated phenotype is likely a consequence of disrupted ciliary signaling. Collectively, these data support an evolutionarily conserved genetic interaction between *Bbs5* and *Nphp4* alleles that may contribute to the variability in ciliopathy phenotypes.

## Introduction

Cilia are highly specialized microtubule-based structures that are evolutionarily conserved from protozoa to primates (Carvalho-Santos *et al.* 2011; Sung and Leroux 2013). While there are exceptions, cilia can be described as either motile or non-motile and are made up of either a 9+2 or 9+0 radially symmetric microtubule scaffold, respectively (Porter 1955). Motile cilia are typically present in multiples on epithelial cells lining surfaces in tissues such as the lungs and reproductive tract and are responsible for movement of viscous materials along the epithelial surface. Non-motile cilia, or primary cilia, serve as a sensory and signaling hub for the cell and in vertebrates, are present on nearly every cell type (Satir and Christensen 2007). In humans, the failure of the cilium to form or function properly manifests in a spectrum of syndromes collectively termed *ciliopathies*. The phenotypes observed among ciliopathies are highly variable (Reiter and Leroux 2017). Classic ciliopathies such as ALMS (Alstrom Syndrome, OMIM #203800) (Tsang *et al.* 2018), BBS (Bardet-Biedl Syndrome, OMIM #209900) (Suspitsin and Imyanitov 2016), JBTS (Joubert Syndrome, OMIM #213300) (Valente *et al.* 2013), MKS (Meckel-Gruber Syndrome, OMIM #249000) (Hartill *et al.* 2017), NPHP (Nephronophthisis, OMIM #256100) (Luo and Tao 2018), OFD (Oral-facial-digital syndrome, OMIM #311200) (Franco and Thauvin-Robinet 2016), PKD (Polycystic Kidney Disease, OMIM #173900) (Yoder 2007), and SLSN (Senior-LØken syndrome, OMIM #266900) (Ronquillo *et al.* 2012) present with pathologies that include, but are not limited to, developmental delay, obesity, hypogonadism, polydactyly, kidney cysts, and retinopathies. Patients can display a wide variability in symptoms with little correlation to specific genetic mutations. One possible explanation for the variability in the phenotypes could be modifier alleles in other ciliopathy genes in the patient’s genetic background.

Following ciliary assembly, a functional primary cilium requires the precise orchestration of multiple complexes to carry out cell signaling events. The cilium’s transition zone (TZ), located at the base, is ideally positioned to function as a barrier between the ciliary compartment and the rest of the cell (Chih *et al.* 2011; Garcia-Gonzalo *et al.* 2011). The TZ is made up of 3 main modules: the NPHP, MKS, and CEP290 complexes (Sang *et al.* 2011). For the cilium to function properly, the TZ not only serves as a barrier but it must also facilitate the passage of necessary materials into and out of the cilium. To do this it must work coordinately with Intraflagellar Transport (IFT) complexes and the BBSome (Zhao and Malicki 2011; Goetz *et al.* 2017; Ye *et al.* 2018). While retrograde (IFT-A) and anterograde (IFT-B) IFT particles are required for ciliary assembly, maintenance, cargo transport, and disassembly, the BBSome is an octameric complex responsible for shuttling transmembrane proteins through the TZ into and out of the cilium but does not typically affect cilia assembly (Nachury *et al.* 2007; Loktev *et al.* 2008; Nakayama and Katoh 2018).

The highly conserved nature of the cilium and its underlying mechanisms allows for studies to be performed in a wide variety of model organisms. Zebrafish have become a powerful tool to better understand the underlying mechanisms of ciliopathies (Drummond 2009). Zebrafish have both motile and non-motile cilia as seen in mammalian systems. Additionally, *ex utero* development and the transparency of zebrafish embryos allow for easy visualization of developmental abnormalities in cilia mutants. *C. elegans* is another powerful tool to study ciliary proteins and to understand the genetic interactions between mutations in ciliary genes. Mutations in cilia genes that would otherwise be lethal in vertebrate models result in easily quantifiable behavioral abnormalities such as altered chemotaxis, defects in dauer formation, and osmotic avoidance defects, but not lethality. *C. elegans* contain a single ciliated cell type, the sensory neuron (Inglis *et al.* 2007). In hermaphrodites, this includes 60 ciliated sensory neurons. A subset of these ciliated neurons, the eight pairs of bilaterally symmetric amphid neurons at the nose of the animal (ASE, ASG, ASH, ASI, ASJ, ASK, ADF, ADL) and 2 pairs of symmetrical phasmid neurons in the tail (PHA and PHB), project through the cuticle exposing them to the external environment (Ward *et al.* 1975). This allows cilia integrity to be readily assayed through a lipophilic dye-filling protocol (Perkins *et al.* 1986). Additionally, the short life cycle of *C. elegans* make them a tractable tool to perform forward genetic screens (Perkins *et al.* 1986; Yee *et al.* 2015).

In previous work, we conducted a modifier screen in *C. elegans nphp-4* mutants to uncover novel alleles that exacerbate ciliopathy phenotypes (Masyukova *et al*. 2016). In support of previous studies showing a genetic interaction between *nphp-4* and *bbs-5* (Yee *et al.* 2015), we identified a novel, and more severe, allele of *bbs-*5. In this work, we extend the previous studies and identify behavioral effects of loss of *bbs-5* and *nphp-4* and demonstrate this new allele is more detrimental than the previously published *bbs-5* deletion allele. To determine if this genetic interaction is evolutionarily conserved, *Nphp4* and *Bbs5* mutant zebrafish and mice were analyzed. Interestingly, double mutant fish do not show an additive effect over the abnormalities shown in *Bbs5* single mutant fish, although assessing possible genetic interactions in the fish model is complicated due to the absence of any overt phenotype in *Nphp4* single mutants. In contrast to the zebrafish model, double mutant mice are nonviable and highlight neurological manifestations that are not detected in either of the single mutants alone. Additionally, we observed reductions in the number of mice containing any combination of three *Nphp4* and *Bbs5* mutant alleles supporting the conservation of genetic interactions between *Bbs5* and *Nphp4* alleles across species. The conserved genetic interactions we observed in these mouse models demonstrate that the overall genetic mutational load in cilia related genes will impact the severity and variability of phenotypes presented by ciliopathy patients.

## Materials and methods

### C. elegans strains

Nematodes were cultivated on NGM agar plates with *E. coli* OP50 bacteria according to standard techniques (Brenner 1974). Nematode culture and observations were performed at 20°C, unless otherwise indicated. The strains used in this study are described in Supplementary Table 1.

### C. elegans Genetic Crosses

All double mutant and reporter strains were generated using standard genetic techniques (Brenner 1974). Homozygosity of alleles and presence of transgenes was confirmed by PCR genotyping and observation of reporter expression using epifluorescence microscopy.

### Generation of Transgenic C. elegans Strains

To generate the strains YH2108 and YH2114, a mixture of the following plasmids was injected into the syncytial gonad of YH1158 or YH1126 adult animals, respectively: 5 ng/µl p341 [*rpi-2p::RPI-2::GFP*], 5 ng/µl p350 [*osm-5p::TRAM-1::tdTomato*], 50 ng/µl pRF4 [*rol-6(su1006)*], 90 ng/µl pBluescript II.

### Dye-filling in C. elegans

Dye-filling assays were performed as described previously with modifications (Perkins *et al.* 1986). Briefly, synchronized L4 animals were washed off NGM plates using M9 buffer and collected into 1.5 ml tubes. Following two washes with M9 buffer, animals were resuspended in 200 µl of M9 and 1 µl of 2 mg/ml DiI (Molecular Probes, Carlsbad, CA) in dimethylformamide (DMF) was added. Animals were incubated in DiI in M9 for 2 hours, rocking gently. After incubation, animals were washed twice with M9 buffer and returned to a fresh NGM plate with OP50. Dye-filling in amphid and phasmid neurons was observed in adults 24 hours later using a Nikon SMZ18 fluorescence stereomicroscope. Worms were scored as “Normal” if all amphid/phasmid neurons showed complete dye-filling, “Partial Dyf” if some dye-filling was lost in the amphids/phasmids, and “Dyf” if there was no dye-filling detected in the amphids/phasmids. For fluorescent imaging, animals were anesthetized using 10 mM levamisole in M9 and immobilized on a 2% agar pad. Confocal images were captured on a Nikon Spinning-disk confocal microscope with Yokogawa X1 disk, using Hamamatsu flash4 sCMOS camera. 60x apo-TIRF (NA=1.49). Images were processed using Nikon Elements and ImageJ software.

### Behavioral Assays in C. elegans

Chemotaxis assays were performed as described (Bargmann 1993; Lee and Portman 2007). Briefly, to prepare assay plates, 8 g of agar was added to 500 ml of deionized (DI) H_2_O, the solution was boiled to fully dissolve the agar, and the following solutions were added after cooling to below 65°C: 2.5 ml 1 M Phosphate buffer, 500 µl 1 M CaCl_2_, 500 µl 1 M MgSO_4_. 10 ml of agar solution was added to 6 cm petri dishes and left to dry for between 3 and 7 days (longer during humid seasons). Prior to the start of assays, plates were marked with spots 4 cm apart to indicate the site of odorant and control (95% ethanol) compounds. Odorant compounds were prepared fresh each time in 95% ethanol at the following concentrations: diacetyl 1:1000; pyrazine 10 mg/ml; 2,3-pentanedione 1:1000; benzaldehyde 1:200. To start assays, worms were collected into 1.5 ml tubes with M9 buffer, then washed twice with M9, and washed once with DI H_2_O. 1 µl of 0.8 M sodium azide was added to marked spots on plates. Worms in a small amount of water were transferred to assay plates at a location equidistant between marked spots. 1 µl each of odorant or control compound was added to the respective marked spots, then a kimwipe was used to wick away excess water from worms. The total number of worms per plate was counted immediately following water removal (50-150 worms/plate). After 60 minutes, the number of worms immobilized at each marked spot was scored. A chemotaxis index (CI) was calculated by subtracting the number of worms at the control spot from the number of worms at the odorant spot, then dividing by the total number of worms counted at the start of the experiment. Each odorant compound was tested at least five times per strain.

Dauer formation assays were performed as described previously (Starich *et al.* 1995). Worms were grown on NGM plates to starvation (∼9 days). Worms were then collected with M9 buffer, spun down, resuspended in 1% SDS in M9, and rocked gently for 1 hour at room temperature. After SDS treatment, worms were washed once with M9 buffer, then transferred to a fresh NGM plate with OP50. The presence of surviving dauer animals was assessed immediately and confirmed after 24 hours.

Egg laying and brood sizes were measured using the following methods. Synchronized L4 animals were singled out on NGM plates with OP50 and allowed to lay eggs for 24 hours. After 24 hours, the adult animals were transferred to fresh NGM plates and the number of eggs and hatchlings laid in the previous 24 hours were counted. This process was repeated every 24 hours until animals from all strains were no longer laying fertilized eggs (4 days). The total number of eggs laid by each individual animal was calculated to determine the brood size. In addition, images were taken at 24 hours post-L4 to show eggs laid during the first 24-hour window of the time course.

To count the number of eggs present in the uterus, synchronized adults at 24 hours post-L4 were transferred to chilled NGM plates. A drop of bleach was placed on each worm to dissolve the cuticle. Remaining fertilized eggs were counted. Eggs were counted from 20 separate animals for each strain.

### Fluorescence Recovery after Photobleaching (FRAP) in C. elegans

To assess the mobility of ODR-10 in the AWA neurons, animals expressing ODR-10::GFP were anesthetized using 10 mM levamisole and immobilized on a 2% agar pad. A FRAP 100 mW 405 nm laser was used to photobleach a portion of the AWA neuron and recovery was observed for 5 minutes.

### Vertebrate Animal Studies

All vertebrate animal studies were conducted in compliance with the National Institutes of Health *Guide for the Care and Use of Laboratory Animals* and approved by the Institutional Animal Care and Use Committee at the University of Alabama at Birmingham.

### Zebrafish Lines

Zebrafish lines were maintained as previously described (Westerfield 2000). The wild-type strain used was AB.

### Generation of mutant zebrafish lines

Alt-R crRNA target sites were designed with Integrated DNA Technologies Alt-R CRISPR HDR Design Tool (https://www.idtdna.com/pages/tools/alt-r-crispr-hdr-design-tool). Alt-R CRISPR-Cas9 crRNA, tracrRNA (IDT, 1072532) and Alt-R S.p. Cas9 Nuclease V3 (IDT, 1081058) was prepared following manufacturer’s instruction. 3 µM sgRNA were obtained through diluting 100 µM crRNA and 100 µM tracrRNA into Nuclease-Free Duplex Buffer (IDT 11-05-01-03), heating at 98°C for 5 min, then cooling to room temperature. The total sgRNA concentration was the same when one or three guides were used. 0.5 µL Cas9 protein was diluted with Cas9 working buffer (20 mM HEPES; 150 mM KCl, pH7.5) to yield a working concentration of 0.5 µg/µL. The diluted Cas9 protein working solution was mixed 1:1 with 3 µM sgRNA solution and then incubated at 37 °C for 10 min to obtain RNP complex. Microinjection was performed by injecting ∼1 nL of RNP complex into yolk of 1-cell stage embryos. RNP complex was fresh prepared and left on ice until microinjection.

### Genotyping with High Resolution Melt (HRM) Analysis in Zebrafish

To isolate genomic DNA from adults, tail clippings from each fish were incubated at 98°C for 20 min in 40 µl 25 mM NaOH in a 96-well plate; then neutralized with 40 µl of 40 mM Tris-HCl. Early-stage or stained embryos were incubated at 55°C for 2 h in 25 µl ELB (10 mM Tris pH 8.3, 50 mM KCl, 0.3% Tween 20, 0.3% NP40, 1 mg/ml Proteinase K) in 96-well plates; then incubated at 95°C for 15 min to inactivate the Proteinase K. PCR reactions contained 1 µl of LC Green Plus Melting Dye (Biofire Defense, BCHM-ASY-0005), 1 µl of 10x enzyme buffer, 0.2 µl of dNTP Mixture (10 mM each), 0.3 µl of MgCl_2_, 0.3 µl of each primer (10 µM), 1 µl of genomic DNA, 0.05 µl of Genscript Taq (E00101), and water up to 10 µl. The PCR reaction protocol was 98°C for 30 sec, then 45 cycles of 98°C for 10 sec, 59°C for 20 sec, and 72°C for 15 sec, followed by 95°C for 30 sec and then rapid cooling to 4°C. Following PCR, melting curves were generated and analyzed using the LightScanner instrument (Idaho Technology) over a 65-95°C range.

### Micro-computed tomography (µCT) on Zebrafish

µCT imaging was performed using the Scanco µCT 40 at a resolution of 16 µm voxels. Contrast enhancement was achieved using Lugol’s iodine solution (Sigma, L6146-1L).

### Zebrafish Histology

Zebrafish were paraffin embedded and histological analyses were performed as described previously (LaBonty *et al.* 2017). The fixation procedure was modified such that 6-month-old adult zebrafish were fixed in 4% paraformaldehyde overnight at room temperature.

### Mice

All animal studies were conducted in compliance with the National Institutes of Health *Guide for the Care and Use of Laboratory Animals* and approved by the Institutional Animal Care and Use Committee at the University of Alabama at Birmingham. Mice were maintained on LabDiet^®^ JL Rat and Mouse/Irr 10F 5LG5 chow.

### Mouse Embryo Isolation

Timed pregnancies were established with embryonic time-point of E0.5 being noted as noon on the morning of observing the copulatory plug. To isolate embryos, pregnant females were anesthetized using isoflurane followed by cervical dislocation. Embryonic tissues or whole embryos were isolated and fixed in 4% paraformaldehyde (Sigma PFA, 158127) in PBS.

### Immunofluorescence microscopy on mouse tissue sections

Ten (10) µm thick tissue cryosections were used for immunofluorescence microscopy. Sections were fixed with 4% PFA for 10 minutes, permeabilized with 0.1% Triton X-100 in PBS for 8 minutes and then blocked in a PBS solution containing 1% BSA, 0.3% TritonX-100, 2% (vol/vol) normal donkey serum and 0.02% sodium azide for one hour at room temperature. Primary antibody incubation was performed in blocking solution overnight at 4°C. Primary antibodies include anti-acetylated α-tubulin (Sigma, T7451) direct conjugated to Alexa 647 (Invitrogen, A20186) and used at 1:1000, anti-Arl13b (Proteintech, 1771-1AP, 1:500). Cryosections were then washed with PBS three times for five minutes at room temperature. Secondary antibodies, donkey anti-rabbit conjugated Alexa Fluor 594 (Invitrogen, 1:1000), diluted in blocking solution were added for one hour at room temperature. Samples were washed in PBS and stained with Hoechst 33258 (Sigma-Aldrich) for five minutes at room temperature. Cover slips were mounted using Immu-Mount (Thermo Fisher Scientific). Fluorescence images were captured on Nikon Spinning-disk confocal microscope with Yokogawa X1 disk, using a Hamamatsu flash4 sCMOS camera. 60x apo-TIRF (NA=1.49) or 20x Plan Flour Multi-immersion (NA=0.8) objectives were used. Images were processed using Nikon’s Elements or Fiji software.

### Tamoxifen Administration in Mice

Recombination of the *Bbs5^flox/flox^* allele was induced in juvenile *CAGG-cre^ERT2^* (Jax Mice stock# 004682) positive mice at postnatal day 7 by a single intraperitoneal (IP) injection of 9 mg tamoxifen (Millipore Sigma, T5648) per 40 g body weight. Tamoxifen was dissolved in sterile corn oil. Adult animals were induced at 8 weeks old by IP injections of 6 mg/40 g body weight tamoxifen, administered once daily for three consecutive days.

### Statistical Analyses

For *C. elegans* chemotaxis and egg laying behavioral assays, we used a two-way ANOVA with multiple comparisons to identify statistically significant data points. For brood size and egg retention counts, we used a one-way ANOVA with multiple comparisons. Analyses were performed with GraphPad Prism 9 (GraphPad Software, LLC, San Diego, CA).

## Results

### Amphid dye-filling is disrupted in C. elegans bbs-5;nphp-4 double mutants

Previously, we conducted a large-scale modifier screen in *C. elegans* to identify genetic interactions able to exacerbate cilia related phenotypes caused by a null mutation in the transition zone component *nphp-4* (Winkelbauer *et al.* 2005; Masyukova *et al.* 2016). From this screen, ten independent strains were isolated, each showing more severe dye-filling defects than *nphp-4(tm925)* mutants alone. Subsequent analysis identified a novel allele in the BBSome complex member, *bbs-5*, in one of the strains from the screen. The *bbs-5(yhw62)* allele is a single nucleotide mutation that results in a tryptophan to stop codon transition in the third exon of the gene (Figure 1A).

**Figure 1.**
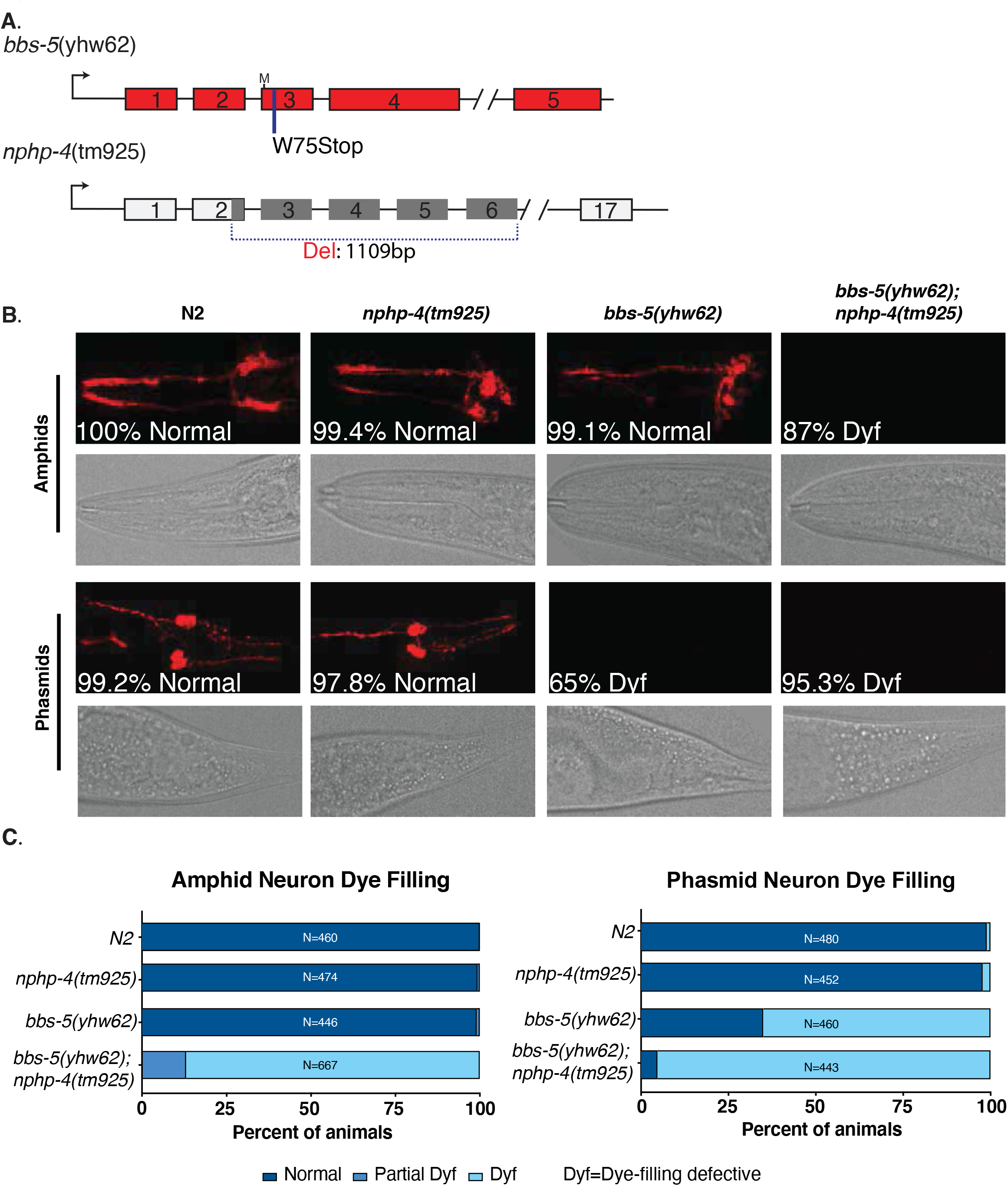
A forward mutagenesis screen identifies genetic interactions between *nphp-4* and *bbs-5* alleles. A) Allele maps of *bbs-5(yhw62)* and *nphp-4(tm925)*. ‘M’ indicates in-frame methionine residue at the beginning of exon 3. B) DiI dye filling of amphid (top) and phasmid (bottom) neurons in *C. elegans.* Percentages indicate frequency of most common dye filling phenotype (see methods for details). C) Quantification of Normal, Partial Dyf, and Dyf phenotypes in amphid (top panels) and phasmid (bottom panels) neurons. N values represent total number of animals analyzed. Dyf, dye filling defective.

Both N2 and *nphp-4(tm925)* worms show normal dye-filling in both the amphid and phasmid sensory neurons (Figure 1B**, quantified in 1C and 1D**). Similarly, the *bbs-5* mutant line obtained from the screen, *bbs-5(yhw62)*, displayed normal dye-filling in the amphid neurons. In contrast, phasmid neurons in most (65%) *bbs-5(yhw62)* animals were dye-filling defective (Dyf). When the *nphp-4(tm925)* mutation was combined with *bbs-5(yhw62)* it resulted in significant dye-filling defects, with 87% and 95.3% of animals showing dye-filling defects in the amphids and phasmids, respectively. Collectively, these data provide support for a genetic interaction between *nphp-4* and *bbs-5* in *C. elegans*.

Additional alleles in the *bbs-5* gene, *bbs-5(gk537)* and *bbs-5(gk507)* have previously been reported, and we compared them with the new allele obtained from the screen. Both the *bbs-5(gk537)* and *bbs-5(gk507)* alleles contain large deletions spanning the putative promoter, the 5’ untranslated region (5’UTR), and the first and second exons (Supplemental Figure 1A). *bbs-5(gk537)* and *bbs-5(gk507)* single mutant animals exhibit normal dye-filling in both the amphid and phasmid neurons (Supplemental Figure 1B**, quantified in 1C**). Double mutant animals harboring either deletion allele display a partial dye-filling defect of their amphid neurons (60.7% in *bbs-5(gk537);nphp-4(tm925)* and 67% in *bbs-5(gk507);nphp-4(tm925)*). A majority of *bbs-5(gk537);nphp-4(tm925)* animals show dye-filling defects in the phasmid neurons (72.2%), while most of the *bbs-5(gk507);nphp-4(tm925)* animals display normal phasmid dye-filling (55.2%).

The location of the deletions paired with the differences between these alleles and the newly generated *bbs-5(yhw62)* allele led us to investigate whether the *bbs-5(gk507)* and *bbs-5(gk537)* are likely to retain some expression and function, perhaps due to the presence of alternate *bbs-5* transcripts. To determine whether any transcript is being made in the *bbs-5(gk507)* and *bbs-5(gk537)* alleles, despite deletion of all predicted promoter sequences, we designed primers downstream of the deletion region (exon 3 to exon 5). The resulting rtPCR generated distinct bands at the predicted size in *bbs-5(gk507), bbs-5(gk537),* and *bbs-5(yhw62)* alleles (Supplemental Figure 1D), suggesting that there is an alternate transcript of *bbs-5*. Importantly, the *bbs-5(yhw62)* alternate transcript still contains an early stop codon in exon 3 that would yield a very short and likely non-functional protein product. The expression of alternate transcripts in the *bbs-5(gk507)* and *bbs-5(gk537)* mutants likely explains the differences in phenotypes observed and suggests that these alleles are hypomorphic, while the *bbs-5(yhw62)* allele is a likely null mutation.

Since the *bbs-5(gk507)* and *bbs-5(gk537)* alleles include a portion of the 3’UTR of the upstream R01H10.7 gene, we performed rtPCR to determine if a read-through transcript was generated. We only detected a minimal amount of possible readthrough transcription in *bbs-5(gk507),* which is likely degraded due to nonsense mediated decay (Supplemental figure 1D).

### Sensory neuron-mediated behaviors are differentially affected by bbs-5 and nphp-4 mutant alleles

Many *C. elegans* behaviors, including chemosensation, dauer formation, and egg laying, have been connected to the proper development and function of the ciliated sensory neurons (Ward 1973; Albert *et al.* 1981; Golden and Riddle 1982; Bargmann 1993; Winkelbauer *et al.* 2005; Lee and Portman 2007). Given the prominent role of cilia in these behaviors we assessed whether any sensory neuron-mediated behaviors might be impacted by the *bbs-5* or *nphp-4* alleles, and whether any behavioral impairments might be exacerbated by the presence of both alleles. First, we assessed chemotaxis to four different attractive compounds: diacetyl and pyrazine, which are recognized by the AWA neurons, as well as benzaldehyde and 2,3-pentanedione, which are recognized by the AWC neurons (Bargmann 1993). For all four odorants, N2 controls yielded an average chemotaxis index (CI) between 0.74 and 0.86, indicative of animals that can detect and preferentially move towards an attractive compound (Figure 2A). In contrast, both *nphp-4(tm925)* and *bbs-5(yhw62)* worms show a significant deficiency in chemotaxis, with average CI values between 0.36 and 0.48 for all four compounds. We did not see any further chemotaxis deficiency in the *bbs-5(yhw62);nphp-4(tm925)* mutants (CI values between 0.34 and 0.5). This suggests that the single mutant *nphp-4(tm925)* and *bbs-5(yhw62)* animals likely have complete loss of their chemotaxis abilities.

**Figure 2.**
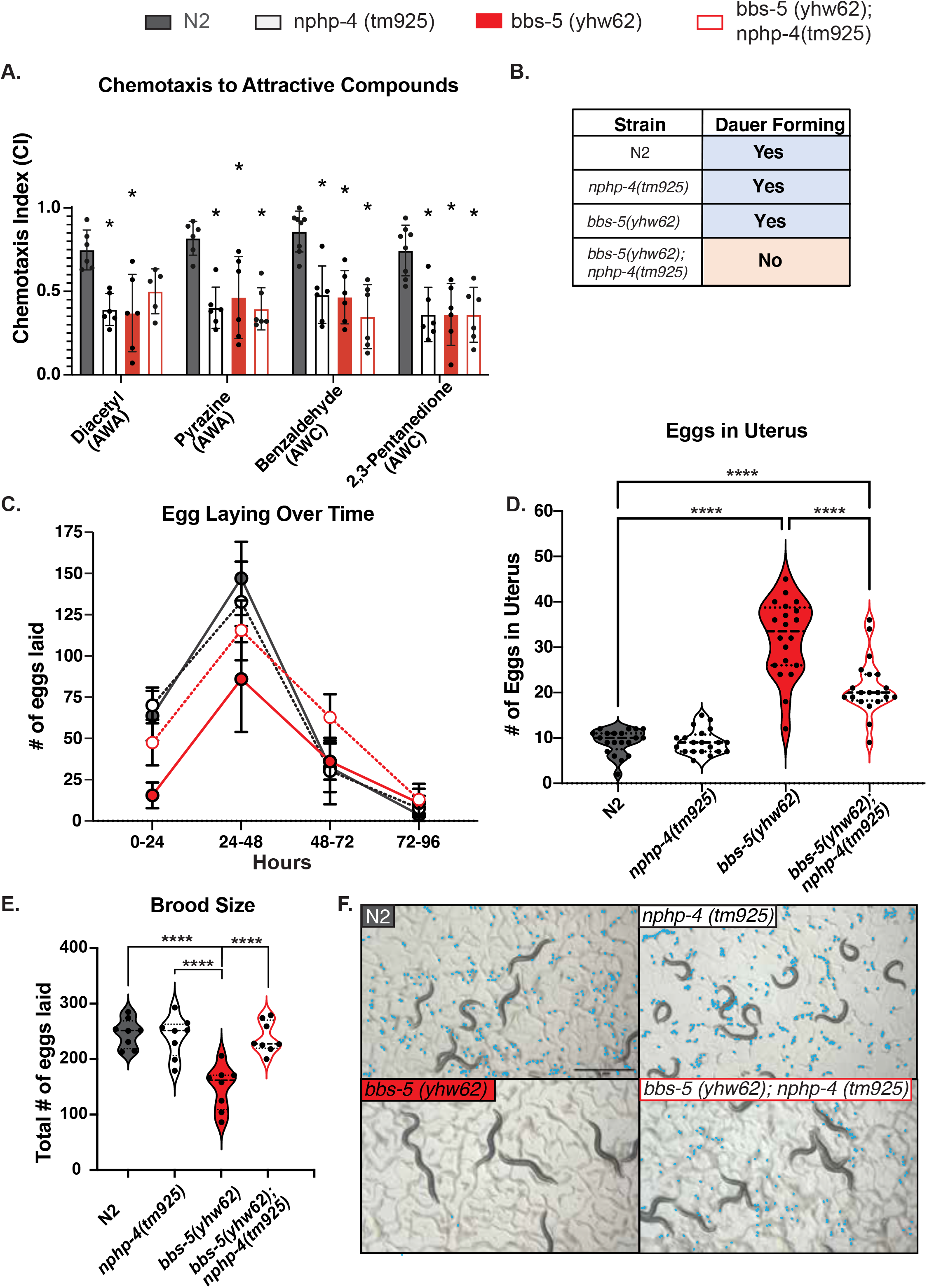
Genetic interaction between *nphp-4* and *bbs-5* alleles alters ciliated sensory neuron-mediated behaviors. A) Chemotaxis to attractive compounds recognized by the AWA neurons (diacetyl and pyrazine) or AWC neurons (benzaldehyde or 2,3-pentanedione) was measured, generating a Chemotaxis index (CI). n ≥50 animals per strain, for at least six replicate experiments. * indicates p<0.05 when compared to N2 control. B) Assessment of ability to form dauer-stage animals. N = 4 replicate experiments. C) Number of eggs laid during 24-hour intervals following L4 stage. n ≥ 6 animals per strain. D) Total number of eggs laid during the course of adulthood. n ≥ 6 animals per strain. **** indicates p<0.0001. E) Visualization of eggs laid in first 24 hours following L4 stage. Each individual egg is pseudo-colored in blue. Scale bar is 1 mm. F) Measurement of eggs retained in the uterus of adults 24 hours following L4 stage. n = 20 animals per strain. Total number of eggs laid during the course of adulthood. n ≥ 6 animals per strain. (Average eggs per strain over time N2: 64 eggs from 48-72 hours, 147 eggs from 72-96 hours, 32 eggs from 96-120 hours, and 4 eggs from 120-144 hours. *nphp-4(tm925):* 70 eggs from 48-72 hours, 133 eggs from 72-96 hours, 30 eggs from 96-120 hours, and 8 eggs from 120-144 hours. *bbs-5(yhw62):* 16 eggs from 48-72 hours, 86 eggs from 72-96 hours, 36 eggs from 96-120 hours, and 11 eggs from 120-144 hours. *nphp-4(tm925);bbs-5(yhw62)* 48 eggs from 48-72 hours, 116 eggs from 72-96 hours, 63 eggs from 96-120 hours, and 13 eggs from 120-144 hours.) *** indicates p<0.001, **** indicates p<0.0001. Error bars in this figure represent standard deviation.

In contrast to the *bbs-5(yhw62)* allele, the two large deletion alleles (*bbs-5(gk507)* and *bbs-5(gk537)*) did not show any significant deficiency in chemotaxis (CI values between 0.7 and 0.79) in the AWA neurons (Supplemental Figure 2A). Interestingly, the *bbs-5(gk537)* mutation was able to rescue the chemotaxis defects caused the *nphp-4(tm925)* allele in double mutants (CI values between 0.62 and 0.72) specifically for AWA neuron detected compounds, but not the AWC-detected odorants. In contrast, the single *bbs-5(gk507)* worms displayed chemotaxis defects specifically to the two AWC-detected odorants, (CI values between 0.27 and 0.62 for benzaldehyde; CI values between 0.22 and 0.71 for 2,3-pentanedione). *bbs-5(gk507);nphp-4(tm925)* double mutants exhibited chemotaxis defects to all four compounds, though CI values ranged widely from 0.16 to 0.82.

Next, we determined whether each strain could enter the dauer stage in response to starvation conditions, another phenotype requiring cilia function (Schafer *et al.* 2006). The N2 control worms and each of the single mutant lines, *nphp-4(tm925), bbs-5(yhw62), bbs-5(gk537)* and *bbs-5(gk507)*, survive SDS treatment, indicating an ability to properly form dauers (Figure 2B, Supplemental Figure 2B). In contrast, all the double mutant lines did not survive SDS treatment, suggesting that the genetic interaction between any *bbs-5* allele and *nphp-4* eliminates the ability to form dauers.

Prompted by the observation that we saw few unhatched, fertilized embryos on plates with *bbs-5(yhw62)* worms (Figure 2E), we analyzed the strains for possible defects in egg laying, egg retention, and total brood size. To address egg laying and total brood size, we age-synchronized hermaphrodites from N2, *bbs-5(yhw62)*, and *nphp-4(tm925);bbs-5(yhw62)* double mutants and counted the number of eggs laid in 24-hour time intervals over four days after the L4 stage. Brood size was determined by summing the total number of eggs laid over the four-day period. The *bbs-5(yhw62)* mutants showed a delay in egg laying over the first 48 hours (Figure 2C, Supplemental Figure 2C) and a significant decrease in total brood size (n=148 eggs, p<0.0001) when compared to the remaining strains, including N2 controls (n=247 eggs, Figure 2D, Supplemental Figure 2D. Surprisingly, the phenotypes observed in *bbs-5(yhw62)* mutants were rescued in *nphp-4(tm925);bbs-5(yhw62)* double mutants. In contrast to the *bbs-5(yhw62)* mutants, brood size in *bbs-5(gk507)* and *bbs-5(gk537)* strains and their corresponding double mutants were not significantly different than controls or *nphp-4(tm925)* single mutants (Supplemental Figure 2D). Although there was some variation across timepoints, overall, none of the strains differed significantly from N2 controls during the final time interval.

We reasoned that the lack of embryos laid in the first 24 hour period and overall delayed egg laying in the *bbs-5(yhw62)* worms could be a result of egg retention (Figure 2E, Supplemental Figure 2E; eggs pseudocolored blue). To test this, we quantified the number of fertilized embryos that were retained in the uterus of age-synchronized young adults. N2 control animals had an average of nine eggs retained in the uterus (Figure 2F). The number of eggs retained by *nphp-4(tm925)*, *bbs-5(gk537)*, *bbs-5(gk507) bbs-5(gk537);nphp-4(tm925)*, and *bbs-5(gk507);nphp-4(tm925)* animals was not significantly different from N2 controls (Figure 2F Supplemental Figure 2F). In contrast, *bbs-5(yhw62)* animals retained more than three times as many eggs, with an average of 31 eggs per animal (p<0.0001). Addition of *nphp-4(tm925)* in the *bbs-5(yhw62)* background partially rescued this defect, with animals retaining an average of 21 eggs in the uterus.

### BBS-5 and NPHP-4 are individually necessary for transition zone integrity

Given that NPHP-4 is a component of the transition zone, which functions as a barrier between the cilium and the rest of the cell, and BBS-5 is a component of the BBSome that mediates both entry and removal of membrane associated proteins from the cilium, we wondered whether there were changes in trafficking of ciliary or non-ciliary proteins in the sensory neurons that may be related to the phenotypic differences observed in the single versus double mutants. We first analyzed RPI-2, a GTPase activator that is normally localized in a region just outside of the primary cilium called the periciliary membrane (Blacque *et al.* 2005; Williams *et al.* 2011) (Figure 3A). When the integrity of the transition zone is disrupted, RPI-2 localization can be seen inside the ciliary compartment (Williams *et al.* 2011). We replicated this result in *nphp-4(tm925)* animals (Figure 3A). Interestingly, we also see RPI-2 expression inside the cilium in both *bbs-5(yhw62)* and *bbs-5(gk537)* mutants analyzed indicating that loss of BBS-5 alone results in improper localization of non-ciliary membrane proteins into the cilium (Figure 3A, Supplemental Figure 3A). Analysis of the double mutants revealed no overt differences in the RPI-2::GFP localization pattern or level of accumulation in the cilium compared to single mutants.

**Figure 3.**
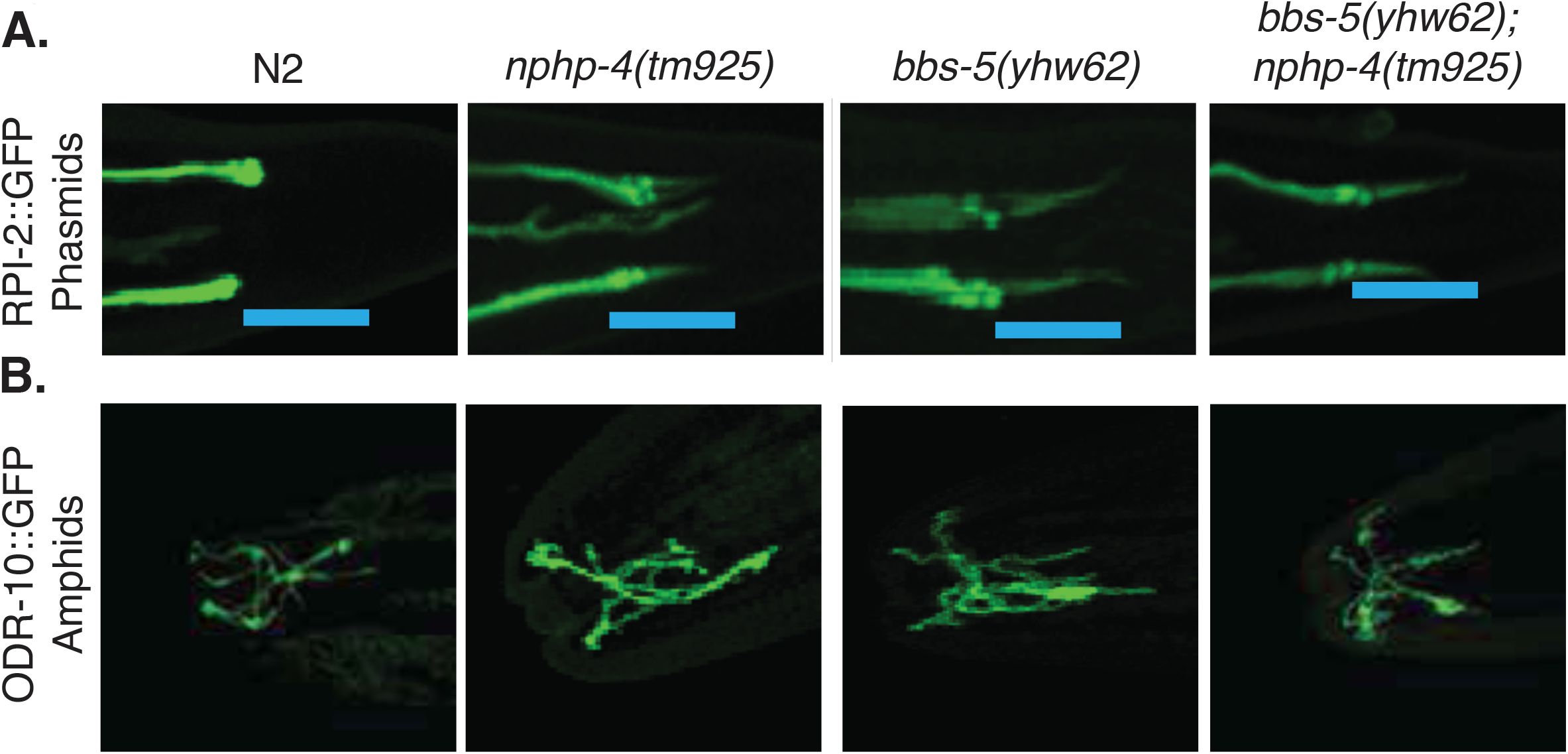
Localization of ciliary ODR-10 and nonciliary RPI-2 membrane proteins is not affected by genetic interaction between *nphp-4* and *bbs-5*. A) RPI-2::GFP protein is excluded from the cilium in N2 worms but enters the cilium in *nphp-4(tm925)*, *bbs-5(yhw62)*, and *bbs-5(yhw62);nphp-4(tm925)* mutants. Data is shown for the phasmid sensilla. B) ODR-10::GFP protein is present in the cilium compartment of AWA amphid neuron of *nphp-4(tm925)*, *bbs-5(yhw62)*, and *bbs-5(yhw62);nphp-4(tm925)* mutants. Representative images selected from among at least 5 animals.

As we saw defects in chemotaxis toward diacetyl in both the *nphp-4(tm925)* and *bbs-5(yhw62)* backgrounds (Figure 2A), we also analyzed whether there are changes in the localization of ODR-10::GFP, the G-protein coupled receptor for diacetyl. ODR-10 is expressed solely in the AWA neurons (Sengupta and Bargmann 1996). In N2 controls, ODR-10::GFP uniformly fills the wing-shaped AWA cilia (Figure 3B). None of the strains carrying mutations in either *nphp-4* or *bbs-5* showed any discernable differences in the localization of ODR-10::GFP (Figure 3B, Supplemental Figure 3B). Furthermore, analysis via Fluorescence Recovery after Photobleaching (FRAP) showed that the trafficking within the cilium is not affected in any of the strains analyzed (Supplemental Figure 4). These data suggest that the chemotaxis defects toward diacetyl observed in the *bbs-5(yhw62)*, *nphp-4(tm925)*, and double mutant strains is not due to loss of the receptor in the cilium but likely a result of disruption of signal transduction downstream of ODR-10.

### Bbs5;Nphp4 double mutant zebrafish do not display exacerbated phenotypes compared to single mutant fish

To investigate whether the genetic interactions between *nphp4* and *bbs5* alleles detected in our *C. elegans* model are evolutionarily conserved, we generated *Nphp4* and *Bbs5* mutant zebrafish using CRISPR/Cas9. The *Nphp4* mutant zebrafish allele (*Nphp4^d7/ d7^*) contains a 7 base pair deletion (TGCACCC) at amino acid 163 (exon 4) resulting in a frameshift. The *Bbs5* mutant zebrafish allele (*Bbs5^+5/+5^*) contains a 6 base pair deletion (CTCTGG) with an insertion of 11 base pairs (AGACAGAGACA) causing a +5 insertion and frameshift at amino acid 8 (exon 1) (Figure 4A). *Bbs5^+5/+5^* mutant zebrafish display severe scoliotic curvature of the spine (Figure 4B). Unexpectedly, no discernable phenotypes were detected in *Nphp4* mutant fish. We then generated *Nphp4^d7/d7^;Bbs5^+5/+5^* double mutant fish. In contrast to the data in *C. elegans*, the addition of the *Nphp4^d7/d7^* mutation did not exacerbate the scoliotic phenotype nor did it present with new phenotypes that are not present in the *Bbs5^+5/+5^* mutant alone. Histological analysis did not reveal gross morphological abnormalities in the vertebrae or vertebral discs in *Bbs5^+5/+5^* and *Nphp4^d7/d7^;Bbs5^+5/+5^* mutant fish (Figure 4C). Similarly, no overt differences were noted by histological analysis of the kidney or heart in *Nphp4^d7/d7^*, *Bbs5^+5/+5^*, and *Nphp4^d7/d7^;Bbs5^+5/+5^* mutant zebrafish (Figure 4C). Histological analysis of the retina indicates that organization of the outer nuclear layer (ONL) is disrupted in *Bbs5^+5/+5^* mutant fish and is not exacerbated in *Nphp4^d7/d7^;Bbs5^+5/+5^* mutant fish (Figure 4C).

**Figure 4.**
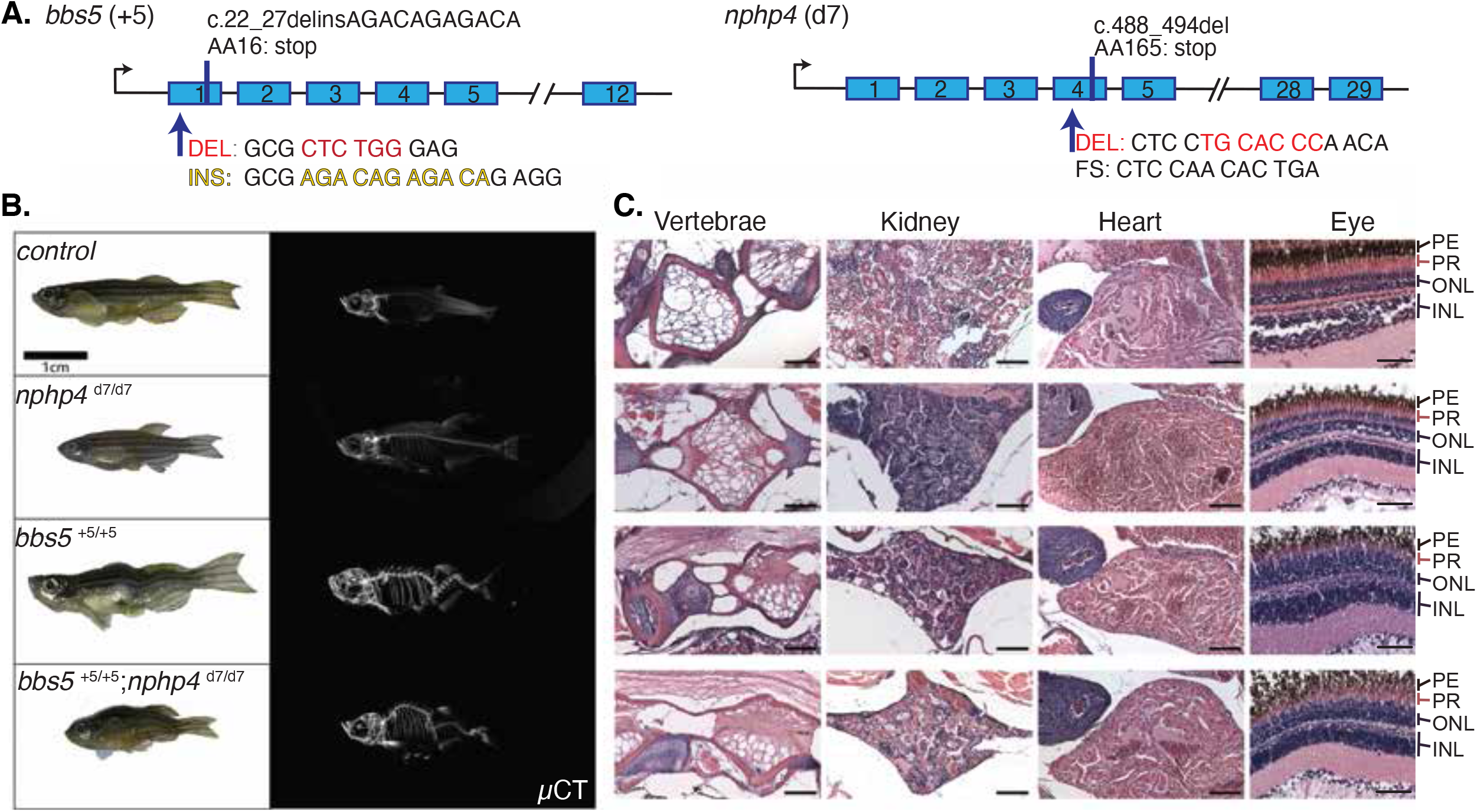
Generation of *bbs5* and *nphp4* mutant zebrafish. A) Allele map of *Bbs5^+5/+5^* and *Nphp4^d7/d7^* mutant zebrafish. B) Micro-Commuted Tomography (uCT) of control, *Nphp4^d7/d7^*, *Bbs5^+5/+5^*, and *Bbs5^+5/+5^*;*Nphp4^d7/d7^* mutant zebrafish. C) H&E analysis of vertebra, kidney, heart and eye in control, *Nphp4^d7/d7^*, *Bbs5^+5/+5^*, and *Bbs5^+5/+5^*; *Nphp4^d7/d7^* mutant zebrafish. Pigmented Epithelium (PE), Photoreceptors (PR), Outer Nuclear Layer (ONL), Inner Nuclear Layer (INL).

### Bbs5^-/-^;Nphp4^-/-^congenital mutant mice do not survive to weaning age

To determine if genetic interactions between *Nphp4* and *Bbs5* mutations are conserved in a mammalian model, we generated *Nphp4*;*Bbs5* double mutant mice (allele maps, Figure 5A and 5B). It has previously been reported that male *Nphp4^-/-^* mice are sterile due to sperm motility defects (Won *et al.* 2011). Similarly, we previously described sterility defects and sub-mendelian ratios at birth and weaning in congenital *Bbs5^-/-^* animals (Bales *et al.* 2020; Bentley-Ford *et al.* 2021). For this reason, double heterozygous *Bbs5^-/+^;Nphp4^-/+^* female by *Bbs5^-/+^;Nphp4^-/+^* male matings were established. Embryos isolated from these crosses show no outward defects at E16.5. Analysis of embryos by contrast-enhanced microcomputed tomography (µCT) similarly showed no obvious defects in tissue morphology including left-right body axis patterning, heart structure, lung, and kidney morphology (Figure 5C). Furthermore, cilia are present in the lateral plate mesoderm of double mutants (Figure 5D). Despite the presence of cilia and the lack of obvious morphological differences between wild-type and double mutant animals, *Bbs5^-/-^;Nphp4^-/-^* mice do not survive until weaning age (Figure 5E) (χ^2^(8, N=103) =37.4, *p*<0.001). Interestingly, there is also a significant decrease in the number of *Bbs5^-/-^; Nphp4^-/+^* and *Bbs5^-/+^;Nphp4^-/-^* animals, further indicating a genetic interaction between the TZ component, *Nphp4*, and BBSome component, *Bbs5* that compromises viability.

**Figure 5.**
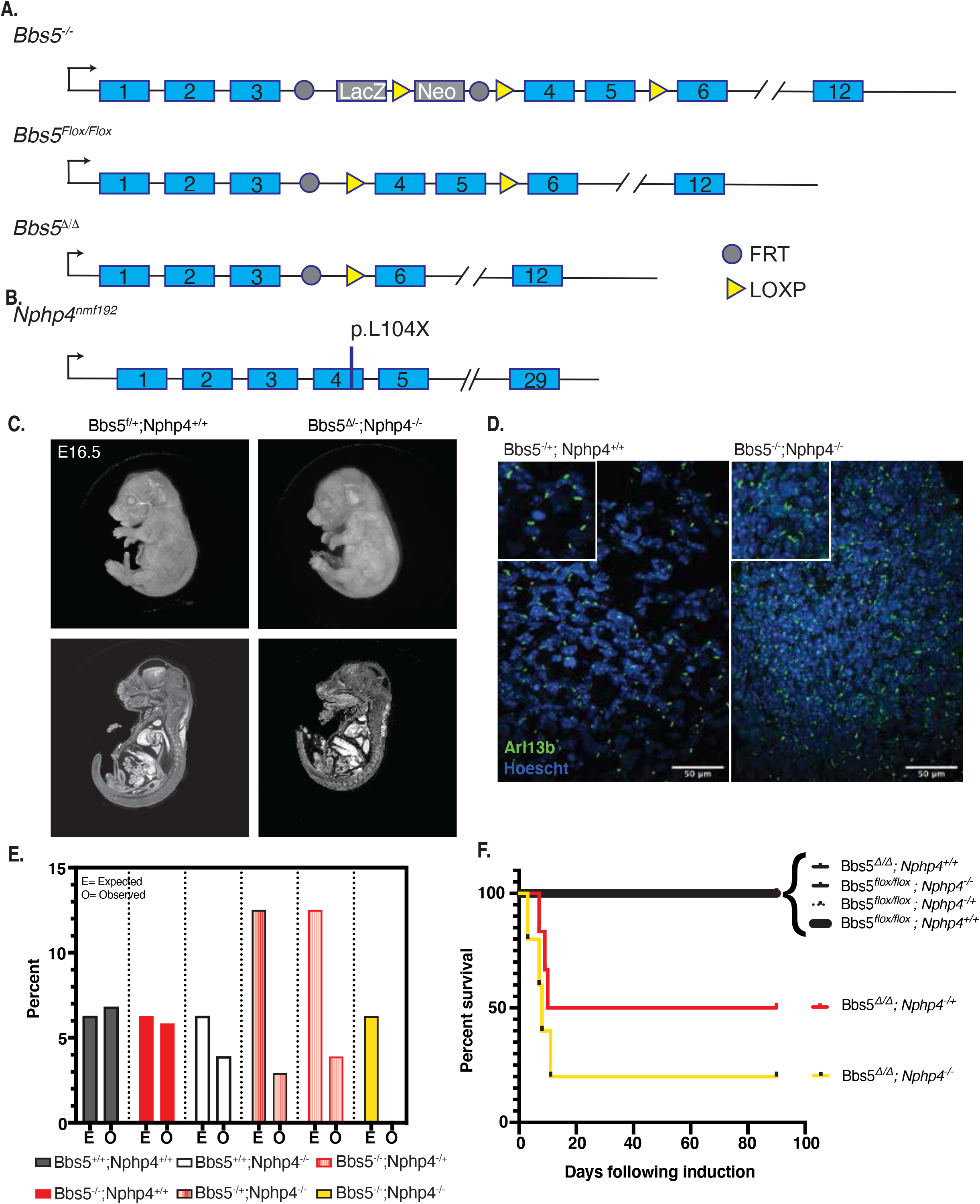
Effect of mutations in *Bbs5* and *Nphp4* mice. A) Allele map of *Bbs5^tm1a^* (*Bbs5^-/-^), Bbs5^tm1c^* (*Bbs5^flox/flox^), Bbs5^tm1d^* (*Bbs5^Δ/Δ^)* B) Schematic depicting the *Nphp4^-^* allele. C) Contrast enhanced micro-commuted tomography of *Bbs5^f/+^;Nphp4^+/+^* and *Bbs5^Δ/-^;Nphp4^-/-^* embryos isolated at E18.5. D) Immunofluorescence staining for cilia (Arl13b, green) in the lateral plate mesoderm of *Bbs5^+/-^;Nphp4^+/+^* and *Bbs5^-/-^;Nphp4^-/-^* double mutant embryos. E) Expected versus observed frequencies of genotypes at weening age generated from a double heterozygous by double heterozygous mating. F) Kaplan-Meier survival curve of P7 induced mice utilizing the conditional *Bbs5* allele (*Bbs5^flox/flox^*) crossed onto *Cagg-Cre^ERT^* wild-type, *Nphp4^-/+^* and *Nphp4^-/-^* mutant backgrounds. When induced at seven days following birth (P7), *Bbs5^Δ/Δ^* (N=7) animals survived at rates comparable to wild-type (N=7) and *Nphp4^-/-^* (N=4) single mutant animals. Comparatively, a significant number of *Bbs5^Δ/Δ^;Nphp4^-/-^* animals (80%, N=5) die following induction. A similar effect is seen in *Bbs5 ^Δ/Δ^;Nphp4^-/+^* animals (50%, N=6), further supporting the existence of a genetic interaction between *Bbs5* and *Nphp4* mutant alleles.

### Loss of BBS5 in a congenital Nphp4^-/-^ mutant background results in neurological abnormalities and lethality

To test for genetic interactions between alleles of *Nphp4* and *Bbs5* postnatally, we utilized the conditional *Bbs5* allele (*Bbs5^flox/flox^*) crossed onto *Cagg-Cre^ERT^*, *Nphp4^-/+^* and *Nphp4^-/-^* mutant backgrounds. Recombination of *Bbs5^f/f^* was induced by IP injection of tamoxifen. When induced at seven days following birth (P7), *Bbs5^Δ/Δ^* (N=7) and *Nphp4^-/-^* (N=4) single mutant animals survived at rates comparable to wild-type (N=7) animals. Comparatively, there is a significant increase in lethality in *Bbs5^Δ/Δ^;Nphp4^-/-^* double mutant animals (80%, N=5) within 20 days following induction. A similar effect is seen in *Bbs5^Δ/Δ^;Nphp4^-/+^* animals (50%, N=6). These data further support a strong genetic interaction between *Bbs5* and *Nphp4* alleles (Figure 5F). *Bbs5^Δ/Δ^;Nphp4^-/-^* animals frequently exhibited uncoordinated behaviors and seizure-like activity (**Supplemental Video 1**). At the time of death, *Bbs5^Δ/Δ^;Nphp4^-/-^* and *Bbs5^Δ/Δ^;Nphp4^-/+^* animals appear runted compared to littermates. The cause of lethality following induction is not known. In contrast to the result obtained with mice induced at P7, animals that are induced at eight weeks of age appear normal and do not express characteristics more severe than either single mutant animals (data not shown). Collectively, these data indicate the genetic interactions between the alleles is important during development and perinatal periods but not in adults.

## Discussion

Using a forward genetic modifier screen, we identify genetic interactions between the ciliary TZ component *nphp-4* and the BBSome component *bbs-5* in *C. elegans.* This interaction was identified via severe dye-filling defects that are present in *bbs-5(yhw62);nphp-4(tm925)* double mutant animals compared to *bbs-5(yhw62)* and *nphp-4(tm925)* single mutants individually. Additional large deletion alleles that include the 5’ end of the *bbs-5* gene, *bbs-5(gk537)* and *bbs-5(gk507)*, were also analyzed. Genetic interactions between *bbs-5(gk507)* and *nphp-4(tm925)* have been previously reported to cause an exacerbated phenotype (Yee *et al.* 2015). However, we show that the *bbs-5(yhw62)* allele identified in our screen displays more severe phenotypes when compared to the *bbs-5(gk537)* and *bbs-5(gk507)* alleles by themselves and when combined with the *nphp-4(tm925)* allele.

The enhanced severity of phenotypes seen with the *bbs-5(yhw62)* allele when compared to the two deletion alleles led us to investigate whether they may retain some function despite the loss of the putative promoter. Our data suggests that both deletion alleles are still able to generate a shorter transcript despite having lost their promoter region. Upon evaluation of the *bbs-5* gene, we identified the presence of an in-frame methionine residue at the beginning of exon 3 (Figure 1A, Supplemental Figure 1A) as well as an SL1 trans-splice leader sequence and several putative transcription factor binding sites suggesting a separate promoter element for generation of an alternative transcript that is predicted to begin at exon 3. These elements are all present in intron 2, which is an uncharacteristically large (1,156 bp) intron for *C. elegans.* The location of the *bbs-5(yhw62)* mutation would still result in an in-frame nonsense mutation within the alternative transcript supporting the conclusion that it is the most severe *bbs-5* mutant allele studied to date, if not the only null allele studied.

Furthermore, when we assayed chemosensing abilities we note that the two large deletion alleles (*bbs-5(gk537)* and *bbs-5(gk507)* are more phenotypically similar to each other compared to the *bbs-5(yhw62)* allele. An exception to this is observed specifically within the AWC neurons. In these neurons we observe chemosensing defects in the *bbs-5(gk507)* mutants alone compared to the *bbs-5(gk537)* mutants. We predict that this variability also stems from the location of the deletions found in these alleles. The transcription factor binding region located in intron 2 includes predicted binding sites for HPL-2, which has been shown to regulate odor adaption specifically in the AWC sensory neurons (Juang *et al.* 2013). The deletion mutation in the *bbs-5(gk507)* allele ablates the entire HPL-2 binding region compared to the *bbs-5(gk537)* deletion, which only disrupts part of the region. This could effectively explain why we see loss of AWC-specific chemotaxis behavior only in the *bbs-5(gk507)* allele. Interestingly, this region is conserved across worms, mice, and humans.

Assays investigating egg laying, the number of eggs retained in the uterus, and brood size also show a defect in *bbs-5(yhw62)* animals alone. Surprisingly, we see a partial rescue of these defects when the *bbs-5(yhw62)* allele is combined with the *nphp-4(tm925)* allele; the mechanisms involved in this partial rescue are not known. Regarding egg laying behaviors, *bbs-5(gk507)* single and *bbs-5(gk507)*;*nphp-4(tm925)* double mutants exhibit moderate delays in the temporal pattern of egg laying. Unlike the results for the *bbs-5(yhw62)* animals, this delay in the *bbs-5(gk507)* animals is not caused by egg retention, nor does it result in a decreased total brood size.

Interestingly, the only behavior that mirrored the exacerbated dye-filling defects seen in all double mutant animals is the ability to form dauers. These results raise a question regarding dye-filling and what it tells us about ciliated sensory neurons. Each of the performed assays are commonly accepted for evaluation of ciliary structure and function in *C. elegans.* The variability seen in double mutant animals between behavioral defects suggests that dye-filling is a readout of a process that is independent of chemosensing ability or transition zone integrity but may be linked to dauer formation. While we do not show disruption of ciliary membrane receptor trafficking based on ODR-10 localization, we do note transition zone integrity abnormalities based on RPI-2::GFP localization. Because these defects are observed in our single mutant animals by themselves, it is not possible to determine whether this is exacerbated in double mutant strains. It is likely that disruption of events downstream of ciliary trafficking in double mutant strains is driving the phenotypes observed.

To determine whether these interactions are conserved across species, *Nphp4* and *Bbs5* mutant zebrafish were generated using CRISPR/Cas9. While *Bbs5* mutant zebrafish exhibit severe scoliotic curvature of the spine and defects in the ONL of the retina, *Nphp4* mutant zebrafish did not exhibit any overt phenotypes. The lack of phenotypes in *Nphp4* fish is surprising considering the effect of mutations in *Nphp4* in mouse models and in human patients, and the important role it has in forming the NPHP complex in the TZ of *C. elegans*. It is possible that *Nphp4* may not be as functionally important in fish or that it preferentially impacts primary cilia compared to motile cilia. Many of the zebrafish ciliopathy phenotypes are associated with defects in motile cilia. Alternatively, it is possible that *Nphp4* mutant zebrafish do develop subtle phenotypes that were simply not detected with the analysis methods employed here or neurological phenotypes that were not analyzed. When combined, *Nphp4;Bbs5* double mutant zebrafish are viable and do not exhibit additional phenotypes when compared to *Bbs5* mutants alone, which is in direct contrast to what we find in the *C. elegans* and mouse models. The lack of observable phenotypes in the *Nphp4* zebrafish makes interpretation of whether a genetic interaction between the alleles is occurring in the double mutant fish challenging.

Previously, we described a congenic mouse model with a mutated *Bbs5* allele (Bentley-Ford *et al.* 2021) and crossed this allele with the *Nphp4* mutant mouse (Won *et al.* 2011) to test whether the genetic interaction occurs in a mammalian model. While mice with either mutation alone are viable, no double mutant animals were obtained at weaning age. More significantly regarding demonstrating a genetic interaction between the alleles, we observed significantly fewer mice that are homozygous for one allele and heterozygous for the other than would be expected. Analysis of embryos at E16.5 revealed that *Nphp4^-/-^;Bbs5^-/-^* mutant embryos appear morphologically normal and that cilia are still present. Thus, the exacerbated phenotypes are not due to defects in ciliogenesis. As we have previously reported, abnormal splicing events in the *Bbs5^-/-^* congenital mouse model causes aberrant tissue specific splicing from the mutant allele, with some tissues able to maintain normal splicing to generate the normal coding region. Due to the complex splicing in the congenital model, we utilized a conditional allele of *Bbs5* crossed with the *Nphp4* allele. Using the inducible Cagg-Cre^ERT^, we were able to induce the loss of BBS5 on the *Nphp4* congenital mutant background. When loss of BBS5 is induced in *Nphp4* mutant juvenile mice (P7), animals frequently exhibit seizures or ataxia, the latter of which could be related to defects in the cerebellum. Induction of BBS5 loss in adult animals (8 weeks old) does not have this effect, supporting a neurodevelopmental consequence caused by combined loss of the *Bbs5* and *Nphp4*. This also points to defects in function of the cerebellum as its maturation occurs during the perinatal period (Chizhikov *et al.* 2007); although we do not observe any overt morphological defects in the cerebellum of the double mutants.

Collectively, our results demonstrate an importance for *bbs-5,* and possibly the BBSome, in interacting with *nphp-4* and the transition zone to regulate ciliary signaling. Future work will aim to identify the disrupted downstream signaling cascades that are responsible for the variety of behavioral phenotypes observed in both *C. elegans* and mice. These data also help address possible mechanisms involved in the phenotypic variability observed in human ciliopathy patients with disease severity being dictated by the overall mutational load in ciliopathy genes that occur in the patients’ genetic backgrounds.

## Data Availability Statement

Strains and plasmids are available upon request. Sequence data is available through GenBank.

## Acknowledgements

We would like to thank the members of Dr. Bradley K. Yoder’s and Dr. John M. Parant’s laboratories for intellectual and technical support on the project. We would like to thank Dr. Maria S. Johnson and Dr. Timothy R. Nagy in the Small Animal Phenotyping sub-core at the University of Alabama at Birmingham for conducting the µCT analysis.

---

**Supplemental video 1. Seizure-like or ataxia activity in P7 induced *Bbs5^Δ/Δ^; Nphp4^-/-^* animals.** *Bbs5 ^Δ/Δ^; Nphp4^-/-^* animal displaying uncoordinated behaviors and seizure-like activity. Bbs5 deletion was induced at P7.

**Supplementary Table 1.**
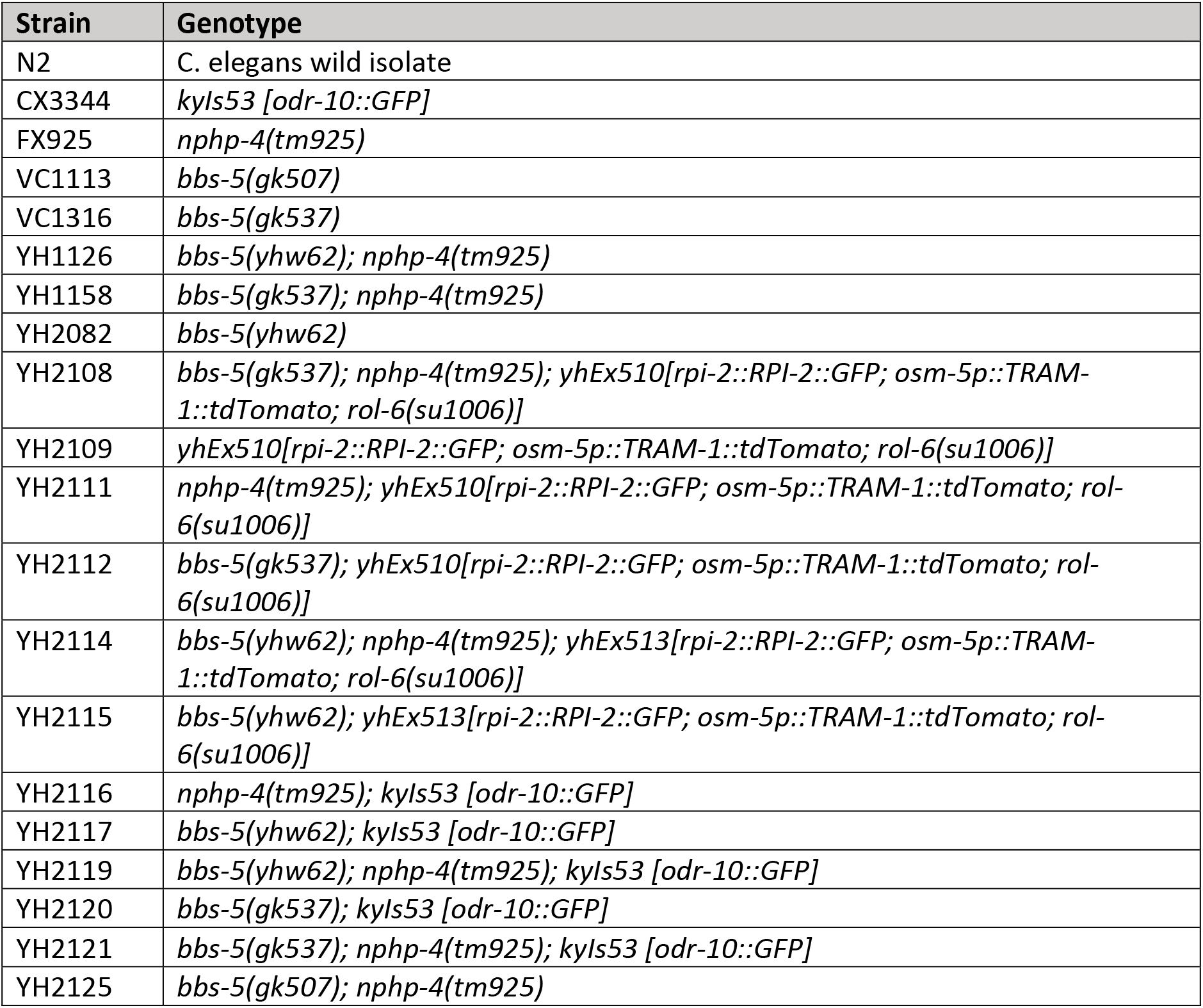
*C. elegans* strains used in studies.

**Supplemental figure 1.**
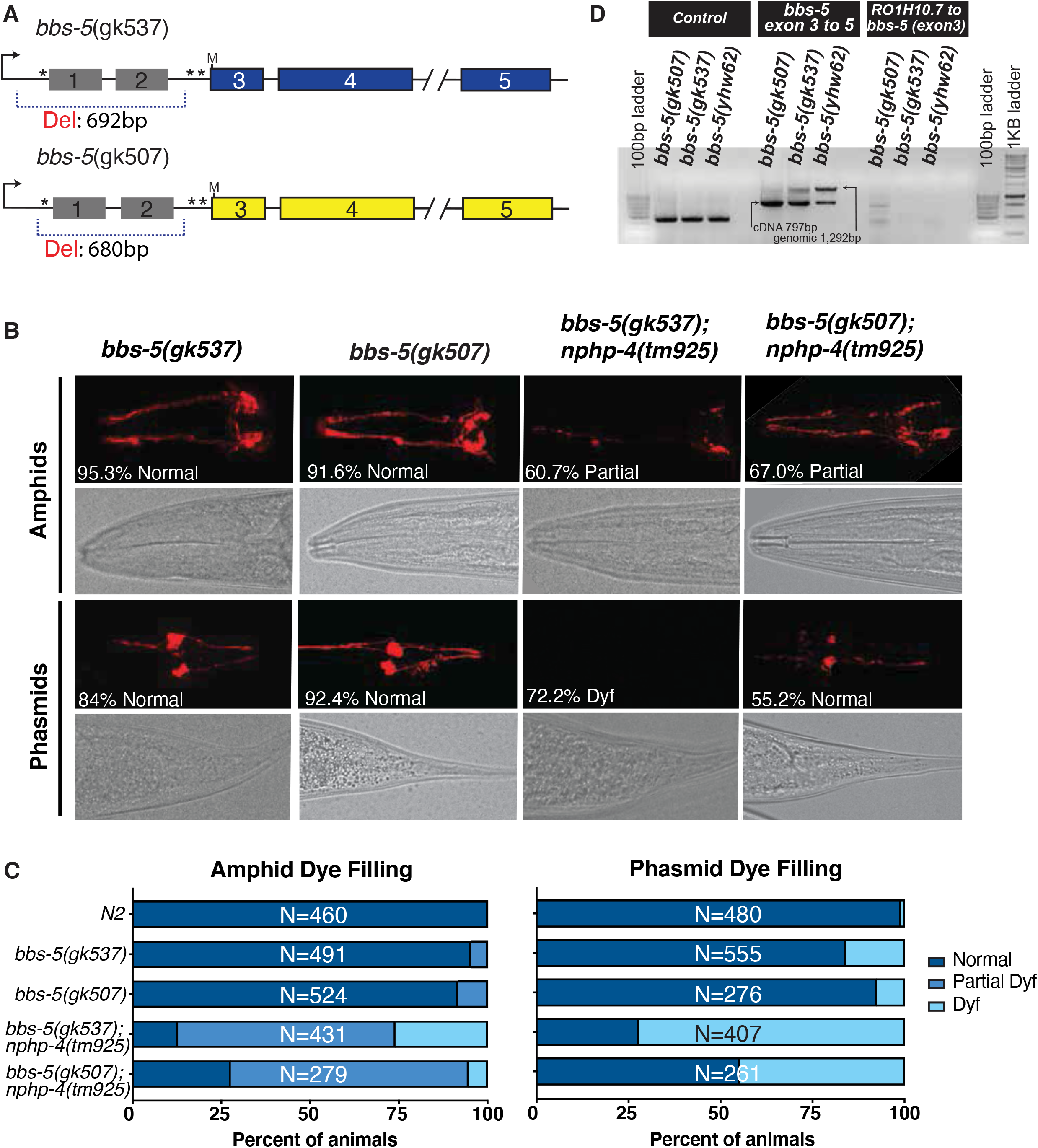
Dye filling analysis of *bbs-5(gk537)* and *bbs-5(gk507)* deletion alleles. A) Allele maps of *bbs-5(gk537),* and *bbs-5(gk507)* with asterisks indicating transcription factor (*) binding sites and ‘M’ indicating an in-frame methionine residue. B) DiI dye filling of amphid (top panels) and phasmid (bottom panels) neurons in *C. elegans.* Percentages indicate frequency of most common dye filling phenotype. C) Quantification of Normal, Partial Dyf, and Dyf phenotypes in amphid (left) and phasmid (right) neurons. N values represent total number of animals analyzed from at least two experiments. Dyf, dye filling defective. D) RT-PCR analysis of transcripts generated from *bbs-5(gk507), bbs-5(gk537)* and *bbs-5(yhw62)* cDNA.

**Supplemental figure 2.**
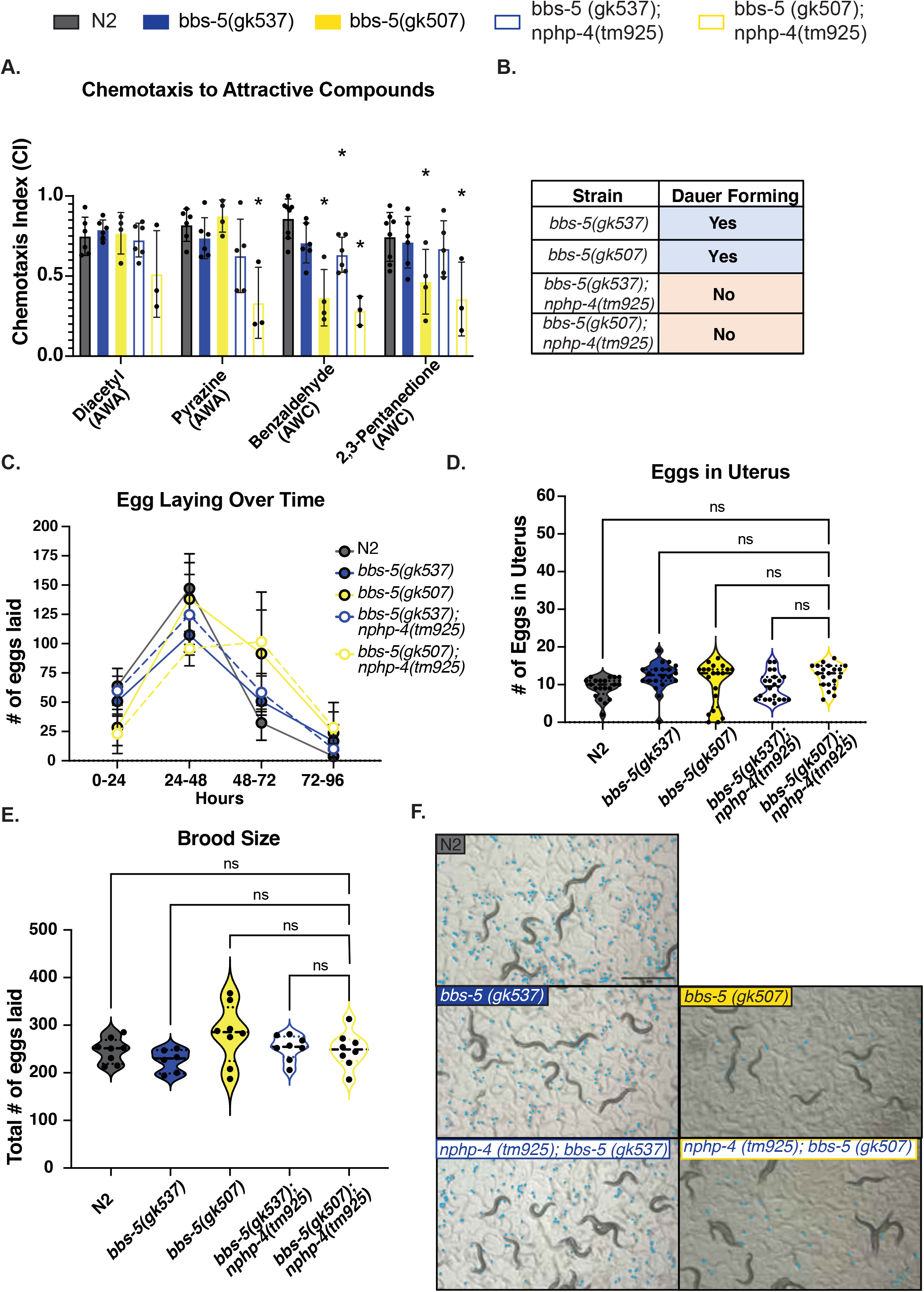
Analysis of sensory neuron mediated behaviors between *nphp-4* and *bbs-5* using *bbs-5* deletion alleles. A) Chemotaxis to attractive compounds recognized by the AWA neurons (diacetyl and pyrazine) or AWC neurons (benzaldehyde or 2,3-pentanedione) was measured, generating a Chemotaxis index (CI). n≥50 animals per strain, for at least three replicate experiments. * indicates p<0.05 when compared to N2 control. B) Assessment of ability to form dauer-stage animals. N = 3 replicate experiments. C) Number of eggs laid during 24-hour intervals following L4 stage. n ≥ 6 animals per strain. D) Total number of eggs laid during the course of adulthood. n ≥6 animals per strain. E) Visualization of eggs laid in first 24 hours following L4 stage. Each individual egg is pseudo-colored in blue. Scale bar is 1 mm. (Average eggs per strain over time *bbs-5(gk507):* 28 eggs from 48-72 hours, 138 eggs from 72-96 hours, 92 eggs from 96-120 hours, and 24 eggs from 120-144 hours. *bbs-5(gk537):* 51 eggs from 48-72 hours, 107 eggs from 72-96 hours, 51 eggs from 96-120 hours, and 17 eggs from 120-144 hours. *nphp-4(tm925);bbs-5(gk507)* 60 eggs from 48-72 hours, 125 eggs from72-96 hours, 58 eggs from 96-120 hours, and 10 eggs from 120-144 hours. *nphp-4(tm925);bbs-5(gk537)* 23 eggs from 48-72 hours, 96 eggs from 72-96 hours, 102 eggs from 96-120 hours, and 28 eggs from 120-144 hours). F) Measurement of eggs retained in the uterus of adults 24 hours following L4 stage. n = 20 animals per strain. ns = not significant. Error bars represent standard deviation.

**Supplemental figure 3.**
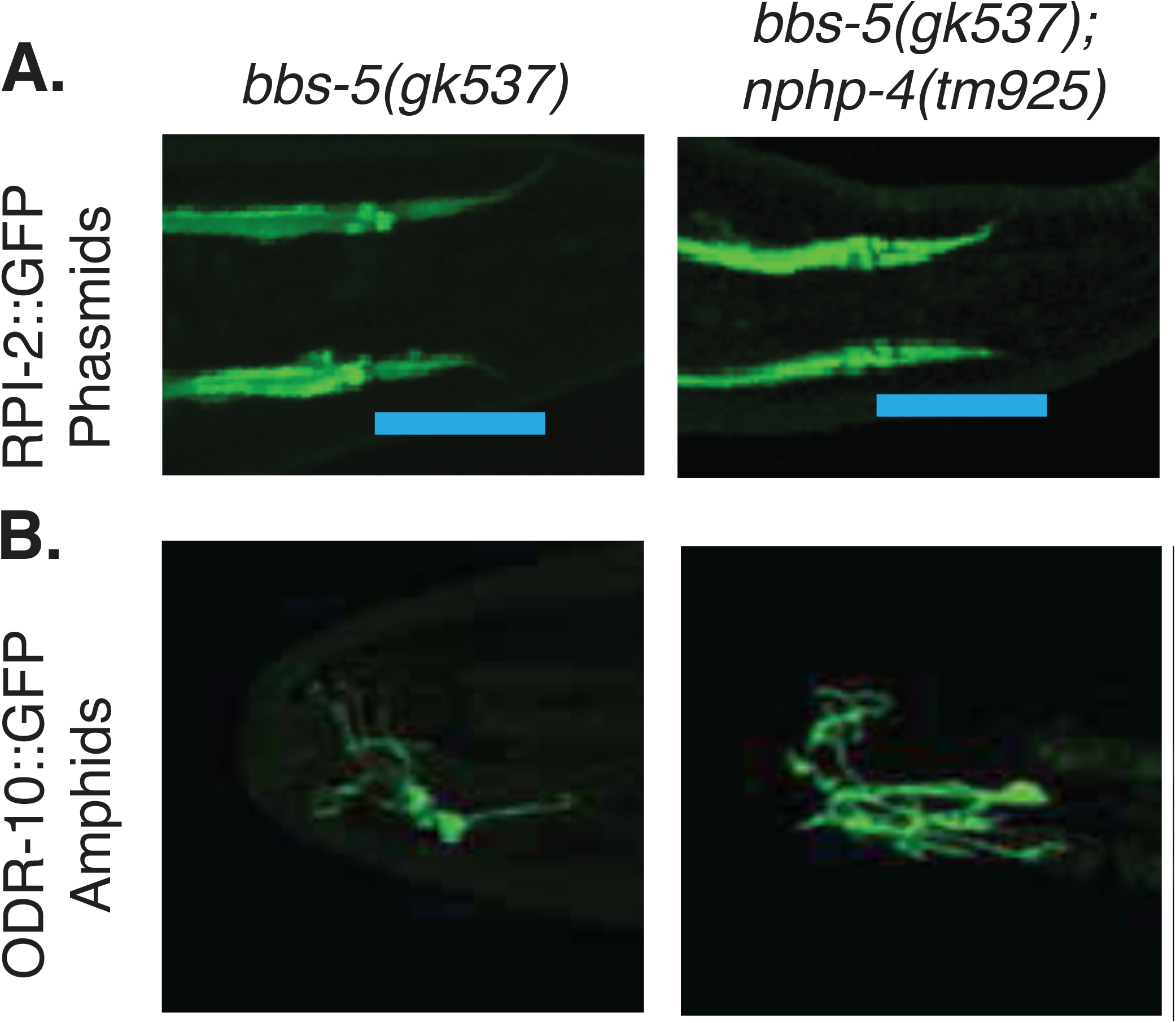
Localization of ciliary ODR-10 and nonciliary RPI-2 membrane proteins in *bbs-5(gk537) and bbs-5(gk537);nphp-4(tm925) double mutants*. A) RPI-2::GFP protein is exclude from the cilium in N2 worms it enters the cilium in *bbs-5(gk537)*, and *bbs-5(gk537);nphp-4(tm925)* double mutants. Data is shown for the phasmid sensilla. B) ODR-10::GFP protein is located normally in the neurons of the amphid sensilla in *bbs-5(gk537)*, and *bbs-5(gk537);nphp-4(tm925)* double mutants. Representative images selected from among at least 5 animals.

**Supplemental figure 4.**
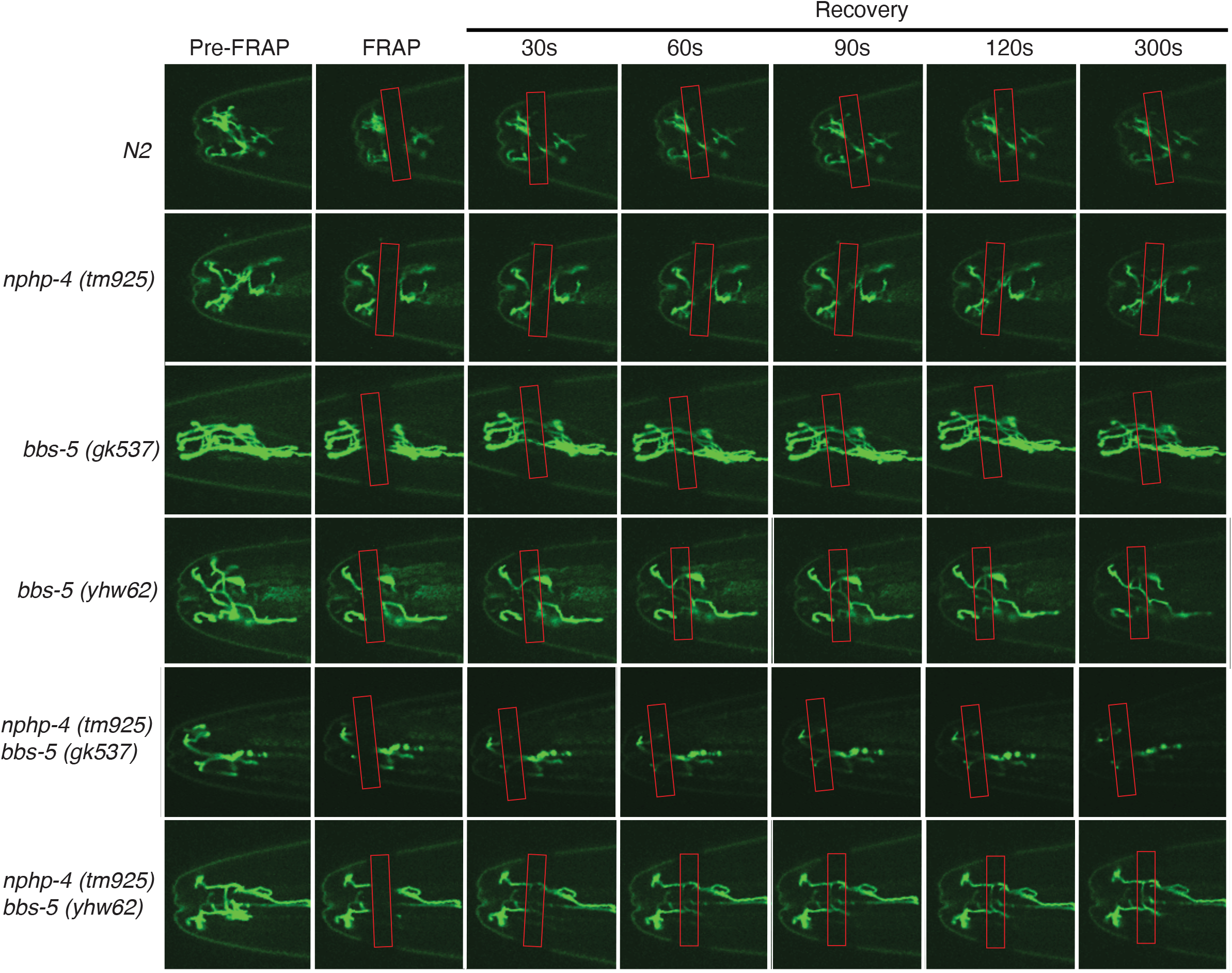
FRAP of ODR-10::GFP reporter. Fluorescence Recovery after Photobleaching (FRAP) of ODR-10::GFP protein in cilia of control and all mutant backgrounds in the neurons of the amphid sensilla over time.

